# Phasing of *de novo* mutations using a scaled-up multiple amplicon long-read sequencing approach

**DOI:** 10.1101/2022.05.06.490885

**Authors:** G.S. Holt, L. Batty, B. Alobaidi, H. Smith, M.S. Oud, L. Ramos, M.J. Xavier, J.A. Veltman

## Abstract

*De novo* mutations (DNMs) play an important role in severe genetic disorders that reduce fitness. To better understand the role of DNMs in disease, it is important to determine the parent-of-origin and timing of the mutational events that give rise to the mutations, especially in sex-specific developmental disorders such as male infertility. However, currently available short-read sequencing approaches are not ideally suited for phasing as this requires long continuous DNA strands that span both the DNM and one or more informative SNPs. To overcome these challenges, we optimised and implemented a multiplexed long-read sequencing approach using the Oxford Nanopore technologies MinION platform. We specifically focused on improving target amplification, integrating long-read sequenced data with high-quality short-read sequence data, and developing an anchored phasing computational method. This approach was able to handle the inherent phasing challenges that arise from long-range target amplification and the normal accumulation of sequencing error associated with long-read sequencing. In total, 77 out of 109 DNMs (71%) were successfully phased and parent-of-origin identified. The majority of phased DNMs were prezygotic (90%), the accuracy of which is highlighted by the average mutant allele frequency of 49.6% and a standard error margin of 0.84%. This study demonstrates the benefits of using an integrated short-read and long-read sequencing approach for large-scale DNM phasing.

## Introduction

*De novo* mutations (DNMs) arise from mutational events that occur during gametogenesis in either parent germ cells, or postzygotically in both somatic and germ cells of the individual carrying them. On average, one to two DNMs can be found in the coding region of a person’s genome (Durbin et al., 2010; O’Roak et al., 2011; Xu et al., 2011). DNMs are of particular significance due to their contribution to many diseases and genetic disorders, notably those affecting individual fitness such as intellectual disability and male infertility (Awadalla et al., 2010; Veltman and Brunner, 2012; Gilissen et al., 2014; Acuna-Hidalgo et al., 2016; Taylor et al., 2019; Oud et al., 2022). It has been shown that approximately 80% of DNMs are of paternal origin (Kong et al., 2012; Goldmann et al., 2016; Yuen et al., 2016; Oud et al., 2022). A major factor known to contribute to an increase in DMNs in individuals is advanced parental age at the time of conception, particularly paternal age (Kong et al., 2012; Goldmann et al., 2016). Investigating the parental origin and timing of DNMs provides not only biological insight into the generation and ability of these DNMs to underlie genetic disorders, it has also been shown to be important for determining the recurrency risk of these disorders (Campbell et al., 2014; Almobarak et al., 2020).

Phasing analysis interrogates the diploid genome, allowing allele separation of the parental chromosomes. This helps not only to determine the parental origin and timing of DNMs, but is also critical to identify compound heterozygous mutations and look into allele specific expression, linked variants, and structural variation (Tewhey et al., 2011; Soifer et al., 2020; Ebert et al., 2021). With short-read whole genome sequencing (WGS) of parent-offspring trios, 15-20% of DNMs can be successfully phased and parent-of-origin called (Goldmann et al., 2016). However, this percentage is expected to be even lower in whole exome sequencing (WES). Phasing challenges can be attributed to the limited sequencing read lengths, the presence of intronic gaps, and the reduced amount of genetic variation in the exonic regions compared to intronic regions (Frigola et al., 2017). By definition, germline DNMs need to be absent in the parental somatic cells, requiring trio-based exome or genome sequencing of parent-offspring trios for discovery. In a next step, the parent-of-origin and zygosity of a DNM can be identified by targeted amplification and long-read sequencing of a region spanning the DNM as well as one or more parentally informative single nucleotide polymorphism (iSNPs). While this appears straightforward, long-read sequencing has both random and positional error, which may result in false variants used for phasing, reducing reliability of downstream analysis (Magi et al., 2018; Watson and Warr, 2019).

There are numerous methodologies to target genomic regions for enrichment prior to sequencing, with the majority being divided into PCR-or CRISPR-based approaches (Hafford-Tear et al., 2019; Gilpatrick et al., 2020; Player et al., 2020). Importantly, when mapping sequence data to the reference genome from CRISPR targeting approaches, the off-target mapping of the sequences is several fold greater than PCR based methods and target coverage is therefore often many factors lower (Hafford-Tear et al., 2019; McDonald et al., 2021), and costs per target are significantly higher. Innate challenges also exist with PCR approaches, including the presence of inhibitors, variable target length, optimisation time, amplification bias, and nucleotide errors (Potapov and Ong, 2017; Shagin et al., 2017). However, despite these challenges with long-range PCR enrichment, the approach is arguably more effective for scaled-up targeted phasing at present.

This study aims to identify the DNM parent-of-origin and zygosity using a targeted long-range PCR approach for phasing 109 distinct DNMs previously identified in infertile men by patient-parent trio exome sequencing (Oud et al., 2022). Targeted amplification of regions encompassing each unique DNM is performed using an optimised long-range PCR workflow designed to quickly increase PCR success rates and reduce pre-sequencing base error. The combination of exome patient-parent trio data, targeted ONT sequencing and validated DNMs are used to improve phasing and allele frequency confidence. Critical aspects of the process are assessed to ascertain practical application for large-scale use of the approach, including amplification length, long-read sequencing error rates and overall phasing performance.

## Methods

### Ethical considerations and study participants consent

77 infertile male patients with unexplained (idiopathic) non-obstructive azoospermia or severe to extreme oligozoospermia and their unaffected parents were enrolled from the Radboudumc outpatient clinic and Newcastle upon Tyne Hospitals NHS Foundation Trust (Newcastle, UK). This study was approved by the ethics Committees/Institutional Review Boards ((Nijmegen: NL50495.091.14 version 5.0, Newcastle: REC Ref: 18/NE/0089). Written informed consent from all patients and their parents was obtained.

### Whole Exome sequencing

Genomic DNA was extracted from whole blood for all patients at the time of evaluation and treatment at the fertility centre. Parental DNA was obtained from saliva using the Oragene OG-500 kit (DNA Genotek, Ottawa, Canada). Exomes of patient-parent trios were prepared for sequencing using 1 µg of high-quality genomic DNA, which was quantified with the Qubit dsDNA HS kit (Thermo Fisher Scientific, Waltham, MA, USA). Whole exome enrichment was performed using Illumina’s Nextera DNA Exome Capture kit (Illumina, San Diego, CA, USA) or Twist Bioscience’s Twist Human Core Exome Kit, as per manufacturer protocols. Individual sample libraries were then indexed with Illumina Nextera DNA UD indexes (Illumina, San Diego, CA, USA) prior to pooling libraries together. The final pooled library was sequenced on the NovaSeq 6000 platform (Illumina, San Diego, CA, USA) to an average coverage depth of 72X (Illumina Nextera kit) or 99X (Twist Bioscience’s kit).

### DNM/iSNP calling and DNM validation

Sequence reads were aligned to the Human Genome Reference Assembly GCRh37 using Burrows-Wheeler Alignment (BWA) version 0.7.17 (Li, 2013) and indexed using SAMtools (Li et al., 2009) and Picard version 1.6 (PicardToolkit, 2019). SNVs and indels were subsequently called by the Genome Analysis Toolkit (GATK)) HaplotypeCaller v4.1.4.1 (McKenna et al., 2010). *De novo* analysis was performed using custom-designed in-house analysis pipelines and validation of DNMs in the proband and parents was performed using standard Sanger Sequencing (Oud et al., 2022). iSNP calling was also performed using an in-house pipeline, which called proband and parent variants using Clair v2 (Luo et al., 2020), proband variants were selected based on unique parental inheritance with a minimum parental coverage of 5x. A single iSNP is selected for parent-of-origin assignment, see section ‘Bioinformatics’

### Target amplification

In 77 infertile males analysed, 109 rare protein coding DNMs were identified and PCR amplified targeted phasing performed, for a basic schematic of the methods see Supplementary Figure 1. Irrespective of known iSNP presence, primers were designed for the target regions (determined as per Figure 1.b) of all 109 DNMs (Supplementary Table 1) using our in-house pipeline which designs primers using Primer3 version 2.3.6 (Untergasser et al., 2012) and Human Genome Reference Assembly GCRh37. Automated rounds of primer creation were run for each target, selecting the most optimal primer pair based on predicted fragment size, self-binding, hairpin factors, reverse primer to forward primer binding, and non-specific genome binding. All expected fragment sizes were limited to a maximum length of 12 kb for greater enrichment success and speed, though optimisation steps were required for 50 % of the targets (Supplementary Figure 2).

**Figure 1.**
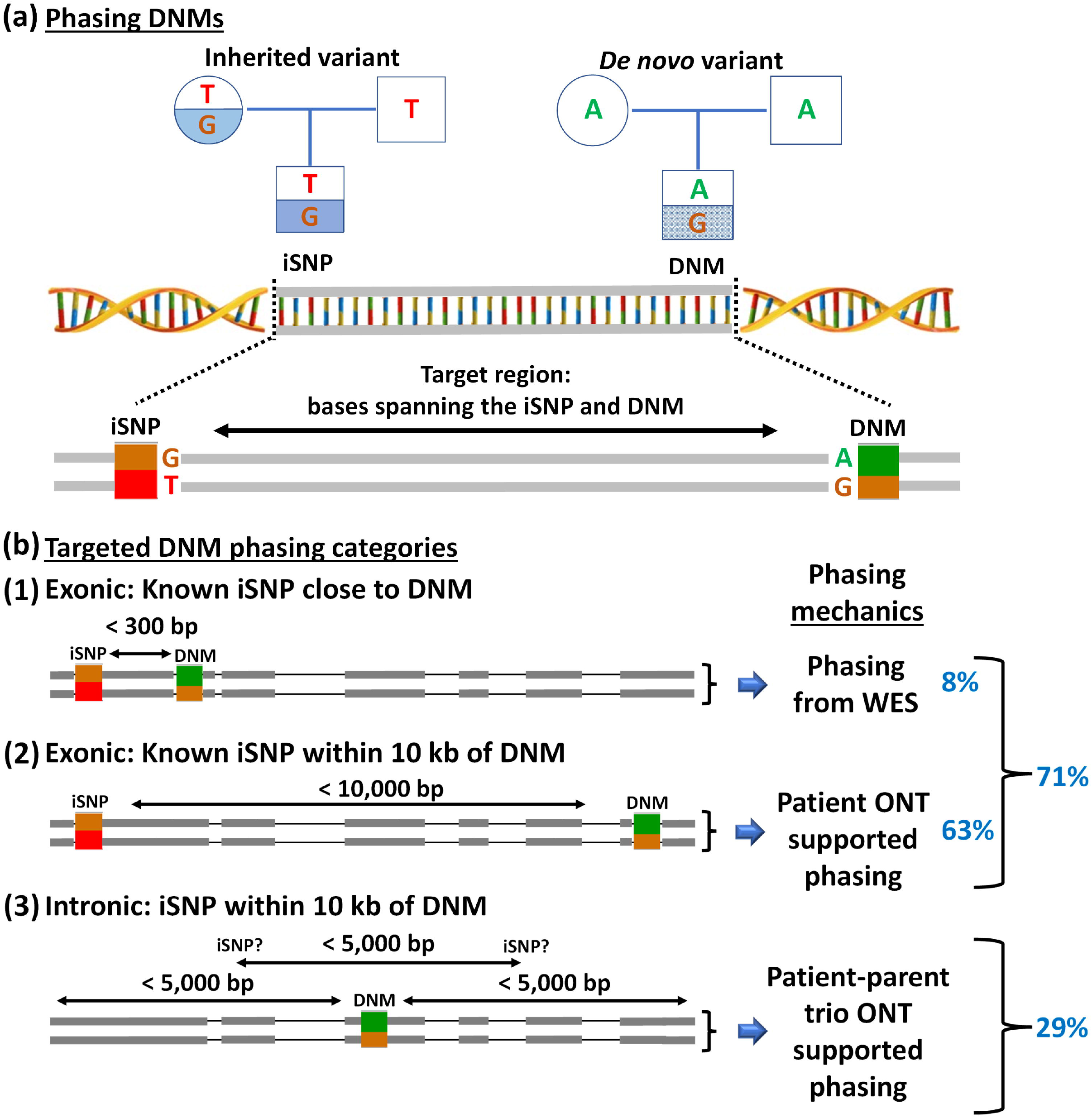
**(a)** Pedigree chart illustrating the bases of an iSNP and a DNM. To phase a DNM and determine the parent-of-origin, single reads must span from the DNM to the iSNP without fragmentation. The distance between these two points of interest dictates the feasibility of phasing the target region with short or long read sequencers. PCR amplification followed by long read sequencing allows target read lengths of up to ∼10,000 bp. **(b)** Illustration of the three categories in which targets are grouped and phased. The grey boxes upon the thin line in each category (1, 2, and 3) represent existing exome read coverage. Each category is based on the distance between the iSNP and DNM, which determines whether it can be phased with only the existing short-read WES trio data (1), WES and long-read proband data (2) or WES and long-read trio data (3). The percentage of samples that fit each category is highlighted in blue.

DNMs without exome supported iSNPs within a 10 kb region were selected for intronic iSNP searching within 5kb (Figure 1.b3). For this group primers were designed in 5 kb windows around the DNM. iSNPs from the ∼5 kb region were identified in the introns of parental ONT data in the downstream bioinformatics. Long-range PCR target enrichment was carried out using optimised running conditions of 2 separate supermixes/enzymes, BioRad iproof and TakaraBio PrimeSTAR GLX (Supplementary Table 2). Primary steps use dilution to reduce contaminants and inhibitors, where dilution was not enough a hotstart supermix (Quantabio RepliQa HiFi ToughMix) was used which reduces the need for annealing temperature specificity (Supplementary Table 2). The number of primer pairs that required bespoke optimisation was reduced to just 12% of the samples by using broad and basic steps as per the Supplementary Figure 2b. Only 5 % of the primer pairs required re-designing, the rest were successfully optimised across the first 5 stages in supplementary Figure 1b. Supplementary Figure 1b steps were chosen based on simple and effective PCR optimisation that could be easily applied on a broad scale for a large number of samples. These measures gave a long-range PCR success rate of >80%, reducing the time required for primer specific optimisation and/or re-design. Sample fragment sizes were confirmed using gel electrophoresis, and quantities were measured with the Qubit dsDNA HS kit (Thermo Fisher Scientific, Waltham, MA, USA), with the best quality supermix enrichment for each given sample/target selected for sequencing.

### Optimising scalability

On average 30 targets per MinION run were amplified, pooled and sequenced. Initially, a limited number of 11 samples were barcoded and sequenced, but a reduction in usable data was noted (Supplementary Table 3). As such, this approach was replaced with careful sample pooling instead. The 11 samples were not re-sequenced, as the small number meant there was still ample data per target (mean of 19 million bases per target). Targets were checked to ensure no overlap in genome location, and demultiplexing was performed based on region extraction of the mapped BAM files (see methods ‘Bioinformatics’). Using this approach, a loss of 17.8% (SEM 7.2) of total basecalled yield was avoided by bypassing barcode ligations, de-multiplexing binning issues and barcode trimming (Supplementary Table 3).

### Long-read sequencing

The long-range PCR target enrichments of >20 ng were prepared for sequencing with the ONT ligation sequencing kit (SQK-LSK109) following the manufacturer’s protocol, with adjustments for sample type and yield. Individual sample libraries with <20 ng were concentrated at given bead clean-up steps. For sample pooling steps, samples were pooled in groups based on fragment size, to allow for more accurate normalisation prior to final pooling and flow cell loading. Prepared samples were sequenced on the MinION using the FLO-MIN106 version 9.4.1 flow cell platform. Flow cells were run until complete pore exhaustion, on average this was 72 hours, with minimal refuel of flow cells performed whenever active pore percentages dropped below 70%, achieving up to 30 billion basecall yields per flow cell and an average coverage depth per sample of >30,000x.

### Bioinformatics: Basecalling, fastq clean up, and demultiplexing

The sequence signal data in multi-fast5 format were basecalled using Guppy version 3.4.4. The resulting fastq outputs were adapter trimmed, low-quality reads ends trimmed (-q 10), and short reads of <30 base pairs removed using cutadapt version 2.5 (Martin, 2011). Cleaned fastq files were mapped against Human Genome Reference Assembly GCRh37 using BWA (Li, 2013) (version 0.7.17), and sample targets were extracted from the resulting BAM file using SAMtools (Li et al., 2009) (version 0.1.19).

### Bioinformatics: Variant calling

Variants were called using Clair v2 (Luo et al., 2020) and filter parameters for variants required >20x coverage, allele frequency 20-80%, 96% confidence, and allele agreement with the validated DNM. The remaining variants were error polished using the high base accurate WES data, removing variants that disagree with any overlapping WES data. WES data for patients and parents was used in filtering ONT called variants illustrated in Figure 2 and Supplementary Tables 4 and 5. Intronic variants identified were further error polished where the inheritance information was available from long-read parental data.

**Figure 2.**
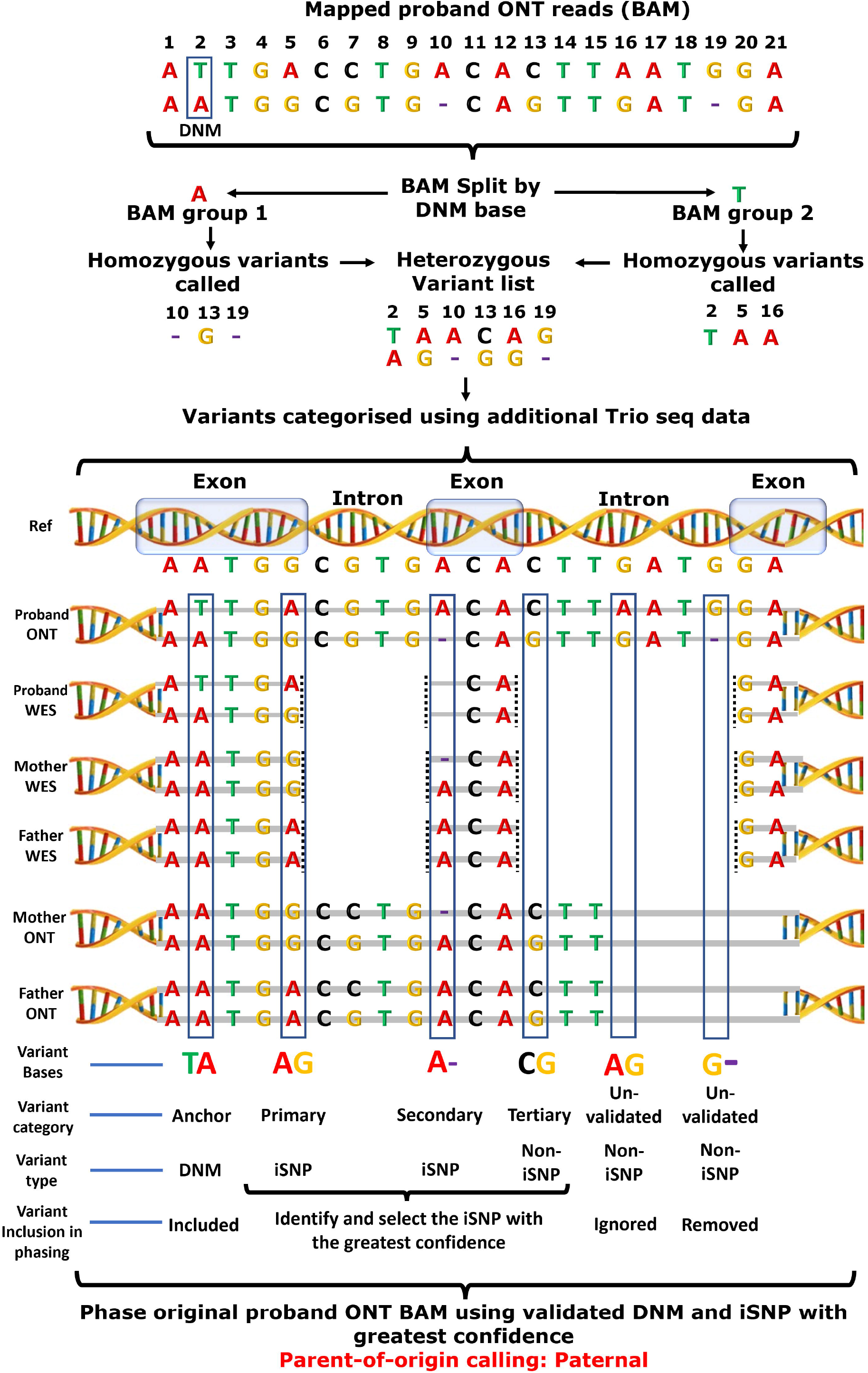
Using validated DNM for an anchored phasing pipeline. Pipeline overview for phasing, including illustration of variant strength and filtering. Reads are split by their Sanger validated DNM nucleotide, then homozygous variants are identified in the two groups. These variants are compared to additional supporting sequencing data and prioritised for phasing based on their primary, secondary, tertiary, and unvalidated categories. These categories are based on the extent of the variant validation from WES and ONT Trio data. Small deletions that have no alternative supporting data are removed due to their high false call rate. An iSNP is selected based on the sequencing support category and used to phase the original proband ONT mapped reads, after which the allele frequencies, zygosity, and parent-of-origin are determined.

### Bioinformatics: Phasing

To identify reliable variants, reads were split based on DNM base type, and the remaining variants were checked for agreement with the split reads using bamql (Masella et al., 2016). The iSNP with the greatest combinations of split read agreement, coverage and additional supporting sequencing data was selected for final phasing, see Supplementary Table 6 for iSNP information. This preliminary variant sorting and iSNP selection approach provided a more reliable variant list. The next step split the raw reads by base type at DNM and selected iSNP positions using bamql. After this step the allele frequencies, parent-of-origin, and timing of the DNM event (pre/post zygosity) was determined. The parent-of-origin conclusion was re-affirmed manually in IGV by visualising the reads containing the DNM and the chosen iSNP from the pipeline.

Additional to manual analysis in IGV, postzygotic and prezygotic DNMs were determined by three category assessments, those that appeared postzygotic are detailed in supplementary Table 11. The categories are; WES DNM base information (coverage/frequency), ONT DNM base information (coverage/frequency), and ONT allele information (coverage and allele frequencies). This information also helped determine background allele error, by presenting reads in the data that represent third and fourth allelic forms.

Allele and base error were calculated in two ways, one approach represented total error, which was calculated for a target DNM by combining all known false base frequencies for total base position error, and for total allele error, combining all known false allele frequencies. The other approach used the most prevalent base or allele frequency that could be determined as false. Assessment of error and noise in data helped support prezygotic and postzygotic DNM calls.

To provide an assessment of basecalling background error that is allele relevant, we ignored checking every base within every target as that would be unnecessarily extensive and time consuming. Instead, we used the highest false base percentage of each iSNP used in phasing each target and calculated the mean false base percentage (Supplementary Table 7). Furthermore, the false base could be qualified as it had parental data support. A quality assessment of the bases for each target within the BAM files was performed using Picard ‘QualityScoreDistributions’ (PicardToolkit, 2019).

## Results

Whole exome sequencing (WES) of 77 infertile males and their unaffected parents identified 109 rare *de novo* mutations (DNMs), all of which were independently validated by Sanger sequencing. Accurate phasing and parent-of-origin calling of the DNMs requires DNA molecules spanning a parentally informative single nucleotide polymorphism (iSNP) and the DNM (Figure 1.a). As such, the ability to call the parent-of-origin is primarily reliant on read lengths. This is notable in the WES data, where only 8% of DNMs could be phased as most iSNP were located >300 bp away from the DNM (Figure 1.b1 and Table 1).

**Table 1.**
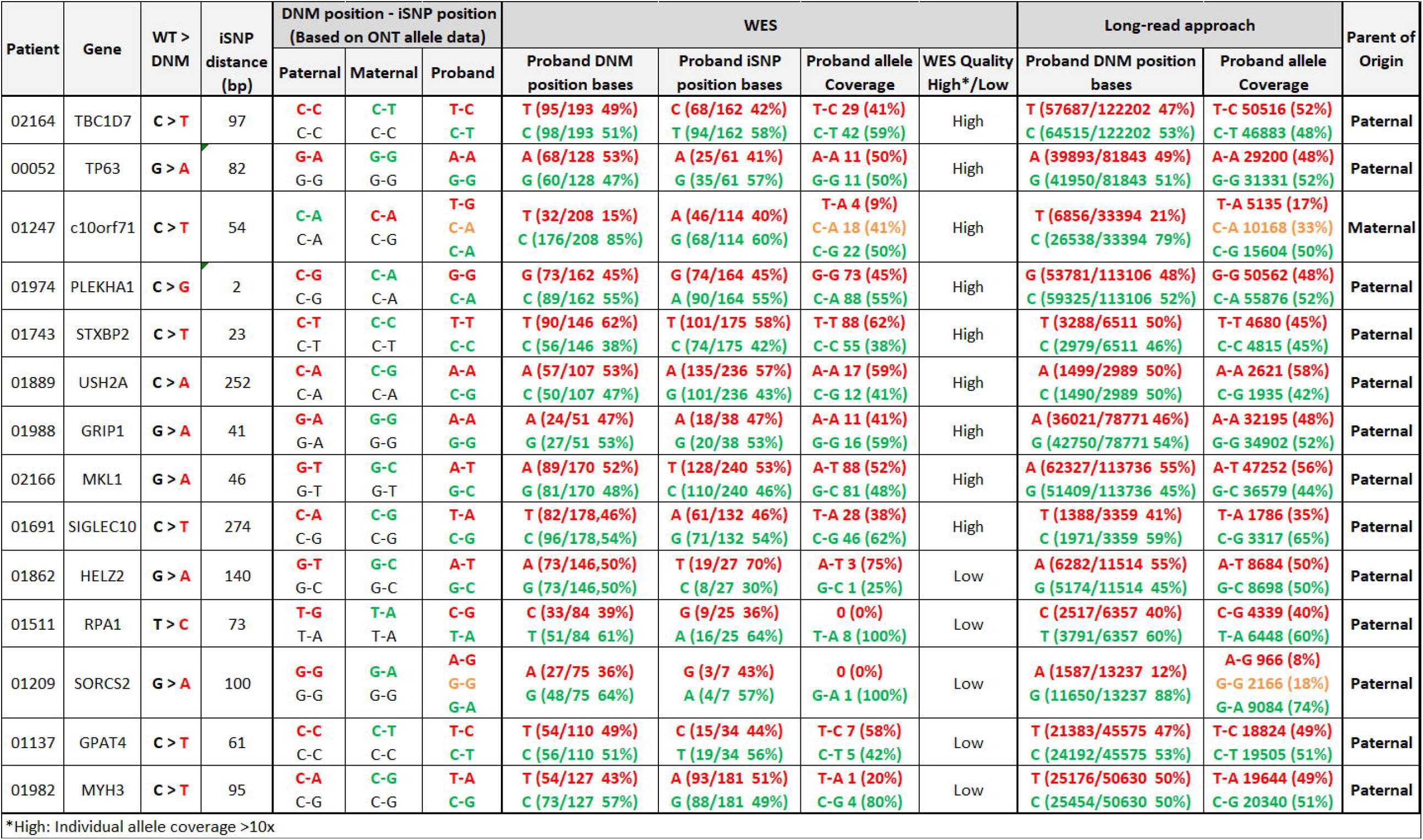
Comparison of WES and targeted long-read phasing. All targets have at least 1 read covering the DNM and iSNP in the WES data. Quality of WES phasing is decerned by the coverage cut-off of >10x coverage per allele. Green bases and alleles represent wt, and red bases and alleles depict the mutant. The given iSNP base pair distance is in relation to the DNM position.

### Target amplification groups

For phasing, all DNM regions were targeted with long-range PCR and sequenced using ONT long-read sequencing. Primer pairs were designed for targeted long-read phasing of the 109 targets (Supplementary Table 1). The fragment size was capped at ∼10 kb to simplify PCR optimisation. Unfortunately, amplification success was still impacted by target length within the 10 kb range, with ideal lengths found to be <4kb (Supplementary Figure 2.a). PCR optimisation steps were required for 50% of all targets (Supplementary Figure 2.b). The amplification size had no impact on error or quality in base calls or allele assignment (Supplementary Figure 3 and Supplementary Table 7). Importantly, for 71% of cases an iSNP was identified in the available trio-based WES data within 10 kb of the DNM (Figure 1.b1 & 1.b2). For this group of DNMs, phasing can be done by targeted long-read sequencing of the proband only, since the iSNP is already typed in patients and their parents.

Cases where an iSNP could not be found in the coding region had primers designed to cover 5 kb regions around the DNM position for parent and proband samples (Figure 1.b3). We chose the 5 kb region based on the analysis of 4344 DNMs identified in 53 whole genome sequenced individuals/children (Smits et al., 2022). This revealed the presence of at least 1 iSNP within 5 kb of any given DNM in 81% of cases (Supplementary Tables 8 and 9). In the end, we obtained long-read sequencing data with iSNPs for 77 out of 109 DNMs selected (71%, Supplementary Table 8 and 10).

### Variants, filtering, error polishing, and phasing

PCR targeted long-read sequencing of the successfully phased 77 DNMs produced an average allele coverage of 35,430X, with a mean background noise of 4% at iSNP positions (see Supplementary Table 7 and methods section ‘Bioinformatics’ for background noise calculation). This increased coverage compared to alternative targeting methods, WES or WGS is expected to help with error reduction and mosaicism detection (Wright et al., 2019). The DNMs, which were initially identified with short-read WES and validated with Sanger sequencing, were used to anchor the preliminary phasing of long reads. This anchoring approach groups long-reads by the base information at the DNM position. After variant calling was performed (methods section ‘Bioinformatics’), homozygous variants were removed and heterozygous variants were checked and filtered based on agreement with the DNM grouped reads. All remaining variants from the long-read sequencing approach were error polished and filtered using WES and parental ONT data (Figure 2 and Supplementary Tables 4 and 5). Following this, iSNPs were identified from the remaining variants. The iSNP with the greatest confidence (coverage, supporting data, DNM allele agreement) was selected for phasing. When phasing the reads based on the DNM and selected iSNP, additional alleles were allowed for the DNM in case of a postzygotic event, but these were screened for credible biological relevance, i.e. the DNM wild type (wt) would have to match the iSNP of the DNM alt. Importantly, this anchored approach filtered out 10% more presumably falsely called variants in comparison to the standard filtering of variants based on quality criteria, sequencing coverage and consensus (see Supplementary Table 4). Small indels made up on average 25% of the false positives, and on average 94% of all indels detected in the long read sequencing data were likely false positives (see Supplementary Table 5, and for an example of a false indel see Supplementary Figure 7). Because of this high error rate for indel calling, we decided to remove all indels without supporting data available, which is noted in the illustration of our approach (Figure 2).

### Validation of the phasing approach using WES data

The parent-of-origin for the DNMs was initially assessed from the short-read WES data. Short-reads, 100bp in length, encompassing DNM and iSNPs in the WES data were available for 14 of the 109 DNMs, but only 9 (8%) (Figure 1.b1) had acceptable coverage (>10x per allele, Table 1). These DNMs were also target amplified and sequenced using ONT long-read sequencing to provide a control cohort for our long-read phasing method. For all 9 DNMs with acceptable coverage in the WES data, the parent-of-origin assignment agreed between the two approaches and the DNM allele frequencies obtained were comparable (Table 1). Interestingly, this was also true for the 5 WES phased samples with limited sequencing coverage, suggesting that the stringent requirement of >10x coverage for WES phasing could be reduced to as little as 4x. For both short and long-read sequencing, 11 out of 14 DNMs showed allele frequencies around 50% (+/-10%), with no significant third allelic form observed (supplementary Table 11), indicating that these were germline DNM events. Of the three DNMs deviating from prezygotic allele frequencies, two were determined as postzygotic (SORCS2 and C10orf71), and one (SIGLEC10) may result from allelic sequencing bias (supplementary Table 12). Coverage of the ONT long-read sequencing for SIGLEC10 was significantly reduced compared to average ONT sequencing coverage, and while both WES and ONT allele data showed DNM allele frequency deviations from 50%, no significant third allelic form was observed. In addition, the DNM base frequencies of both WES and ONT data were within the prezygotic range of 50% +/-10%.

For the DNM affecting the C10orf71 gene in patient 01247, the *de novo* mutated allele (T-A) had a much lower allele frequency of 9% and 17% when detected with both WES and ONT approaches respectively, with similar percentages observed in DNM base frequencies. Apart from the wt allele and the DNM allele, we clearly observed a third allelic form that represented the wt version of the DNM allele. The presence of this third allelic form suggests that the DNM likely occurred as a postzygotic event. The postzygotic mutation is observed in both the WES and ONT data in this case. Postzygotic mutations can, however, be missed when WES has low coverage, as seen for the DNM in SORCS2 in patient 01209. The postzygotic DNM in SORCS2 presents an average discrepancy of 15% from prezygotic norms in the WES base frequencies. A third allelic form is not shown in the WES allele data due to only having 1x coverage. Greater base discrepancies are observed in the ONT base frequencies, with an average deviation from prezygotic norms of 38%, and with several thousand times more coverage in the ONT data, a wt of the DNM allele is observed at 18%.

### WES vs ONT DNM allele frequency comparison and parent-of-origin

In total, 77 DNMs of 109 DNMs (71%) were phased and the parent-of-origin determined, with 64 of these (83%) being of paternal origin. Across the 77 phased DNMs there was a DNM allele frequency range of 6% to 78% in the ONT data and 15% to 71 % in the WES DNM base data, clearly showing the presence of postzygotic and prezygotic mutations. From these 77 DNMs, our targeted long-read sequencing approach identified 61 clearly prezygotic DNMs, with a possible third allelic form found in the remaining 16 DNMs. The separation of pre and postzygotic DNMs can be established more clearly through a combined analysis of their base frequencies, allele frequencies, DNM wt allele frequency, and background sequencing error (see Supplementary Table 11). By doing this we confirmed 69 prezygotic DNMs and 8 postzygotic mutations, as depicted in Supplementary Table 11 and 12, and Supplementary Figure 4 and 5. On average a third of allelic data is identified as false in our ONT approach (Supplementary Table 7), and that impacted the allele frequency significantly. After removal of these false allelic data, we noticed that the average ONT allele frequency for prezygotic DNMs were around 50% with an average of 49.6%, with a standard error margin (SEM) of 0.8% (Supplementary Table 13 and Supplementary Figure 4). For germline DNMs, we expect the mutant allele frequency to be approximately 50%. shows that the majority of all ONT DNM allele frequencies were around 50% with the prezygotic average of 49.6% (SEM 0.8%). While raw allele frequencies were expectedly much worse than DNM base frequencies, the polished allele frequencies are slightly more accurate compared to the WES DNM and ONT DNM prezygotic nucleotide base frequencies of 48.6% (SEM 1.0%) and 48.8% (SEM 0.8%), respectively (Supplementary Table 13). As expected, postzygotic frequencies of mutant alleles deviated significantly from this, with an average of 15.9% and a standard error margin of 2.4% in ONT allele data, though the frequencies of postzygotic mutations is less consistent in WES data, where the postzygotic frequency average was 28.8% (SEM 4.9%) (Supplementary Table 13).

The parent-of-origin was also investigated within the pre and postzygotic fractions, where we observe the prezygotic paternal preference of 83% and the postzygotic paternal origin of 62% (5 out of 8), see Supplementary Figure 5. Classification of postzygotic DNMs was determined as per Supplementary Table 11 and a detailed IGV illustration is depicted in Figure 3. All postzygotic calls were corroborated by the DNM base frequencies in the short-read and long-read data, and further supported by all identified allele frequencies and the degree of error in target data (see Supplementary Table 7 and 11).

**Figure 3.**
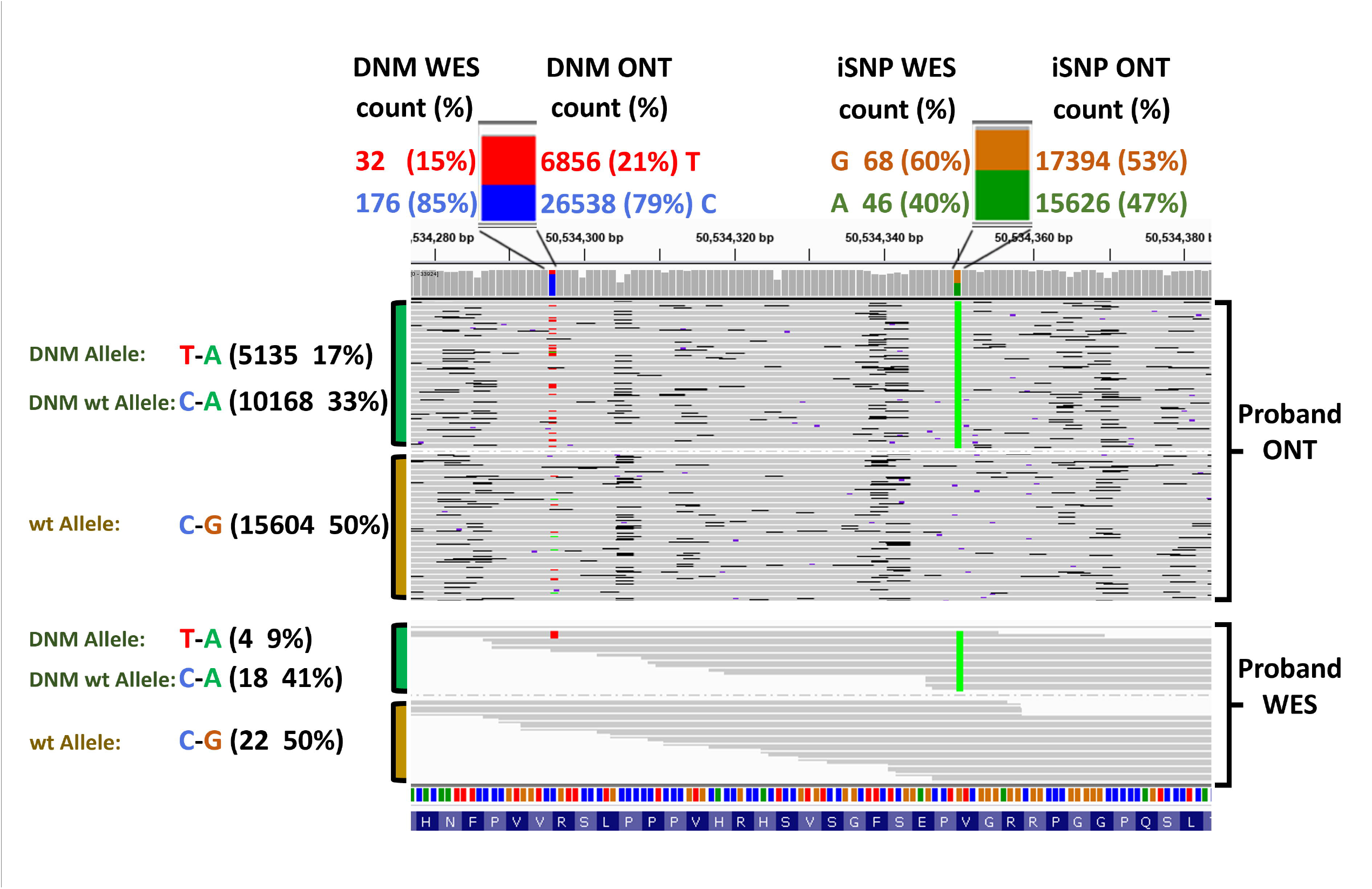
Example of the postzygotic DNM in C10orf71 visualized in IGV. The iSNP and DNM base associated (coloured) counts and percentages are displayed at the top of the plot as a descriptive extension of the coverage bar. This postzygotic DNM is observed in the ONT and WES allele data and is supported by the discrepancies in the DNM positional base frequencies, while the iSNP base frequencies remain close to the expected 50:50 ratio. The IGV visual of the reads presents them grouped by iSNP base type. This read view illustrates a third allelic form in the reads that matches the base and iSNP combination expected, had the DNM not occurred (C-A 33% in ONT, 41% in WES).

## Discussion

In this study we optimized and applied a multiplexed long-read sequencing approach which makes use of high-quality short read exome data to perform routine phasing of *de novo* mutations. We report the phasing results of 77 DNMs from 64 of our 77 patient-parent trios linked to male infertility, achieving successful phasing for 71% of the 109 DNMs investigated using this long-read targeted approach. In contrast, only 9 of these DNMs (8%) could be reliably phased based on short-read WES data alone.

Short-read exome sequencing has become an increasingly common tool in research and diagnostic of genetic disease, with patient-parent trio-based sequencing routine for the detection of DNMs. With only 8% of DNMs being phasable in our cohort of 77 patients when using short-read WES alone, it is clear that an alternative approach is needed to determine the parent-of-origin and timing of DNMs. Our method uses the long-range PCR with standard optimisation steps to achieve the simplest and quickest large-scale success. PCR is a simple and standard wet lab practice, providing greater enrichment and target specificity than any alternative target-based approach. The sequencing of target enriched long DNA strands with ONT allowed us, in most cases, to acquire 10s of thousands of times coverage per target, with many targets run per flow cell, supporting the projects scale demands. To overcome challenges with error and postzygotic mutations we used the WES data and Sanger validated DNMs to polish the variant analysis, which limited computational demand as no complex algorithms were required, and processing could be quick.

DNMs are known to arise from mutational events occurring during gametogenesis, predominantly during spermatogenesis rather than during oogenesis, which is assumed to be associated to the scale of male gamete production and failure of DNA repair mechanisms which lead to the increased opportunity for mutational events to occur (Aitken & Baker, 2020; Evenson et al., 2020; Grégoire et al., 2013; Haldane, 1947; Kong et al., 2012). Previous literature has shown that DNMs occur on the paternal allele approximately 80% of the time (Kong et al., 2012; Goldmann et al., 2016; Yuen et al., 2016). In agreement with this literature, 83% of all phased DNMs in this study were determined to be of paternal origin.

Parent-of-origin and zygosity information adds another layer to our understanding of potential disease-causing variants. This is important when investigating genetic diseases, especially those that likely have complex and varied mechanisms. In our cohort of 77 patients, 51 patients were confirmed to suffer from non-obstructive severe oligospermic or azoospermic phenotypes. In the original publication related to this work (Oud et al., 2022), we showed that 6 out of the 8 likely causative DNMs identified in these patients were of paternal origin (Supplementary Table 8 and 14, Supplementary Figure 6). This suggests that DNMs with a deleterious effect on the health of an individual can escape negative selection in the paternal germline.

Accurate detection of the DNM allele frequency is critical to differentiate prezygotic from postzygotic mutational events, important in clinical settings for estimating the recurrence risk (Almobarak et al., 2020; Scanga et al., 2021). Our approach yielded a highly accurate allele frequency average of 49.6% in the prezygotic mutations, with an SEM of 0.84% (Supplementary Table 13). Though similar accuracy may be achievable with more computationally demanding methods, the strength of our method lies in utilizing the WES data and DNM validation practices commonly available. This shows that bioinformatic cleaning and more complex haplotype processing steps are unnecessary, with accurate results achievable through simple DNM and DNM-anchored iSNP selection. In total, 8 of the 77 phased DNMs were classified as postzygotic events (10%), largely in agreement with current literature results of 6.5% to 10% (Acuna-Hidalgo et al., 2015; Ye et al., 2018; Sasani et al., 2019), supporting the validity of our method. Interestingly, while there was significant correlation between WES and ONT postzygotic base/allele frequencies, 25% of the postzygotic DNMs could not be determined from WES DNM base frequencies. This demonstrates the importance of combining phasing analysis with deep coverage long-read sequencing to further characterise the timing of DNMs. As can be expected for postzygotic DNMs (Girard et al., 2016), we see less paternal bias even though our numbers are small (5 out of 8 postzygotic DNM are paternal, 62%).

We here use a standard PCR amplicon targeting approach with long-read sequencing, rather than CRISPR-Cas targeting. Despite CRISPR-Cas recently becoming a choice method for long-read targeted sequencing (Hafford-Tear et al., 2019; Liu et al., 2019; Gilpatrick et al., 2020; McDonald et al., 2021), the large number of targets and small target sizes in our cohort would make CRISPR-Cas complex and costly. Standard PCR targeting is optimal for routine application that does not require methylation data, read lengths greater than 10-20 kb, or directly representative read counts (Aird et al., 2011). While the CRISPR-Cas approach can have a 10-100 fold enrichment of the target region compared to standard low coverage long-read WGS, it still results in 95.4 % off-target sequencing (Gilpatrick et al., 2020). This off-target sequencing issue significantly limits the number of samples that can be run per flow cell, and only a single sample can be run if demultiplexing is based on the genomic position of the target. The reverse is seen when comparing this to the standard amplicon approach used herein, where dozens of samples were run per flow cell and no off-target mapping was identified. Based on using the optimal CRISPR-Cas approach of 2-3 gRNAs, and taking into account the reduced sample number per flow cell, CRISPR-Cas methods also have >40 fold increase in cost per target.

Nonetheless, CRISPR-Cas target enrichment shows great promise, and will likely be the best approach for targets larger than 10-20 kb. Despite not observing more basecalling error from PCR extension in targets of greater sizes, it is worth considering the potential increases in base error and bias from PCR approaches which would compound the lower accuracy inherent to long-read sequencing. Our data supports the importance of minimizing target region sizes when performing PCR based amplification for targeted sequencing, especially when performing primer optimisation for >100 bespoke primer pairs. Limiting target sizes will reduce labour intensive PCR optimisation and though not observed in our study it may also reduce base error from PCR fidelity issues. We should, however, be mindful that for 11% of the DNMs studied no iSNPs were found within the 5kb window, so minimizing the target region can also negatively impact phasing. For another 18% of DNMs, however, the sequencing data was of insufficient quality for phasing purposes, so clearly a balance must be found between sequencing quality and target size.

Since ONT released the MinION platform in 2014, there have been extensive leaps in advancing both the chemistry and the bioinformatic tools. This has resulted in raw base accuracy moving from as low as ∼60% (Loman and Watson, 2015) to the current 92-97% in the 9.4.1 flow cell chemistry used in this investigation. It should be noted that further increases in accuracy have also been suggested in recent flow cell chemistry, such as the release this year of the R10.4.1 flow cell. Bioinformatic tools that include the variant caller ‘Clair’, used here, have also shown increased confidence in variant calls but are thought to be reaching their limit, with greater confidence requiring significant alternative algorithms or improvements in chemistry (Luo et al., 2020). Despite the bioinformatic improvements in base calling and variant calling, we observe that the accuracy of long-read data on long-range PCR products still causes far greater false positives than WES short-read data. After filtering ONT variants by read depth and quality scores, our anchored approach filtered an additional 50% of the remaining variants on average. If false variants that were missed prior to our anchored filtering approach were included in the phasing process it is likely some targets would be phased incorrectly or not phased at all. Many phasing tools such as ‘whatshap’ carry out phasing with the understanding that variants within the vcf file are correct, so the removal of false variants is important.

Our study provides an approach for accurately phasing and parent-of-origin calling DNMs in a set of 77 patients. To our knowledge this is the first time that phasing of DNMs has been investigated on this scale using long-range PCR targeted ONT sequencing, where each sample has a uniquely specific target. We optimized the method for efficiency and streamlined the laboratory and computational pipelines for processing large numbers of DNMs for detailed phasing analysis. We incorporate additional short-read sequencing patient-parent trio data and Sanger validated DNMs that are commonly available from DNM discovery pipelines like ours. This approach enabled us to improve DNM phasing and postzygotic calling. This data-supported and anchored phasing approach can be of great use in both research and diagnostic settings where DNMs are routinely studied and interpreted.

## Authorship

Study was designed by J.A. Veltman and G.S. Holt. G.S. Holt performed primer design, some primer optimisation, ONT sequencing, bioinformatics, data analysis, manuscript curation. L. Batty performed primer optimisation, ONT sequencing and DNM Sanger validation. B. Alobaidi performed DNA extractions, WES sample prep, some PCR optimising, some ONT sequencing, and DNM Sanger validation. H. Smithperformed DNM selection and disease causative classifications of DNMs. M.S. Oud performed DNM selection, manuscript review and disease causative classification of DNMs. L. Ramos provided clinical data. M.J. Xavier performed DNM identification in WES data and manuscript review. J.A. Veltman manuscript review. All authors contributed to the final paper.

## Acknowledgements

We appreciate the participation of all patients and their parents in this study. We thank Arron Scott and Bryan Hepworth (Newcastle University) for technical support, the genomics core facility for whole exome sequencing (WES), the Bioinformatic support unit (BSU) for initial WES processing. This project was funded by an Investigator Award in Science from the Wellcome Trust (209451) to J.A.V.

## Data Availability Statement

Further data is available upon request from authors

## Supplementary Tables

**Supplementary Table 1** DNM target region primer list.

**Supplementary Table 2** Long-range PCR reaction mixes and running conditions.

**Supplementary Table 3** Base loss breakdown of the 11 barcoded samples after demultiplexing and barcode removal.

**Supplementary Table 4** Point mutation breakdown of different filtering steps.

**Supplementary Table 5** Indel breakdown of different filtering steps.

**Supplementary Table 6** Selected iSNP information for each target.

**Supplementary Table 7** DNM target region breakdown of data error and quality.

**Supplementary Table 8** Data overview. Summary tables of supplementary data.

**Supplementary Table 9** Dataset from Smits et al., 2022 used to calculate the percentage of DNMs that have an iSNP within 5kb ranges.

**Supplementary Table 10** Failed target sequenced samples.

**Supplementary Table 11** Data analysis for determining the post-zygotic DNMs.

**Supplementary Table 12** Phasing and parent-of-origin determination in WES and ONT datasets.

**Supplementary Table 13** Mutant base and allele frequencies with associated pre/post zygotic classifications.

**Supplementary Table 14** DNMs in this dataset with previous causative classifications for male infertility (Oud et al., 2022).

## Supplementary Figures

**Supplementary Figure 1 Basic overview of the long-read targeted phasing approach.** Red boxes highlighting failed sample outcomes.

**Supplementary Figure 2 Long range PCR performance and optimisation. (a)** change standard to rapid. Percentage comparison of first-time primer success using our standardised approach and primer success after optimisations at increasing target sizes (kb). **(b)** Table of optimisation categories, displaying total target amplification success rates at each stage.

**Supplementary Figure 3 a)** Log10 scatter plot of target amplification sizes in relation to the percentage of false allele coverage of each target and the percentage of false iSNP/DNM base coverage of each target. The trend of the data is highlighted by the logarithmic line of best fit. **b)** Log10 scatter plot of target amplification sizes in relation to the percentage of bases with quality score <5 (see method section ‘Bioinformatics’). The trend of the data is highlighted by the logarithmic line of best fit.

**Supplementary Figure 4** Violin plot comparing the WES DNM nucleotide base frequencies, ONT DNM nucleotide base frequencies, ONT DNM allele frequencies, ONT DNM prezygotic and postzygotic allele frequencies.

**Supplementary Figure 5** Pre and post zygotic illustrative breakdown of the 77 DNMs that could be phased, including the parental origin of the DNMs.

**Supplementary Figure 6** Stacked percentage plot looking at parent-of-origin in relation to likelihood of causality.

**Supplementary Figure 7 IGV illustration of a DNM showing a comparison of WES and ONT data.** Using long-read targeted sequencing, the ∼1700 bp region that these variants span was sequenced and visualized in IGV. Approximately 15000 reads are phased in the ONT data, and the quality and coverage differences between those reads and the WES reads can be observed.

