## Supplementary figures and images for "Phasing of *de novo* mutations using a scaled-up multiple amplicon long-read sequencing approach"

### Supplementary Figure 1

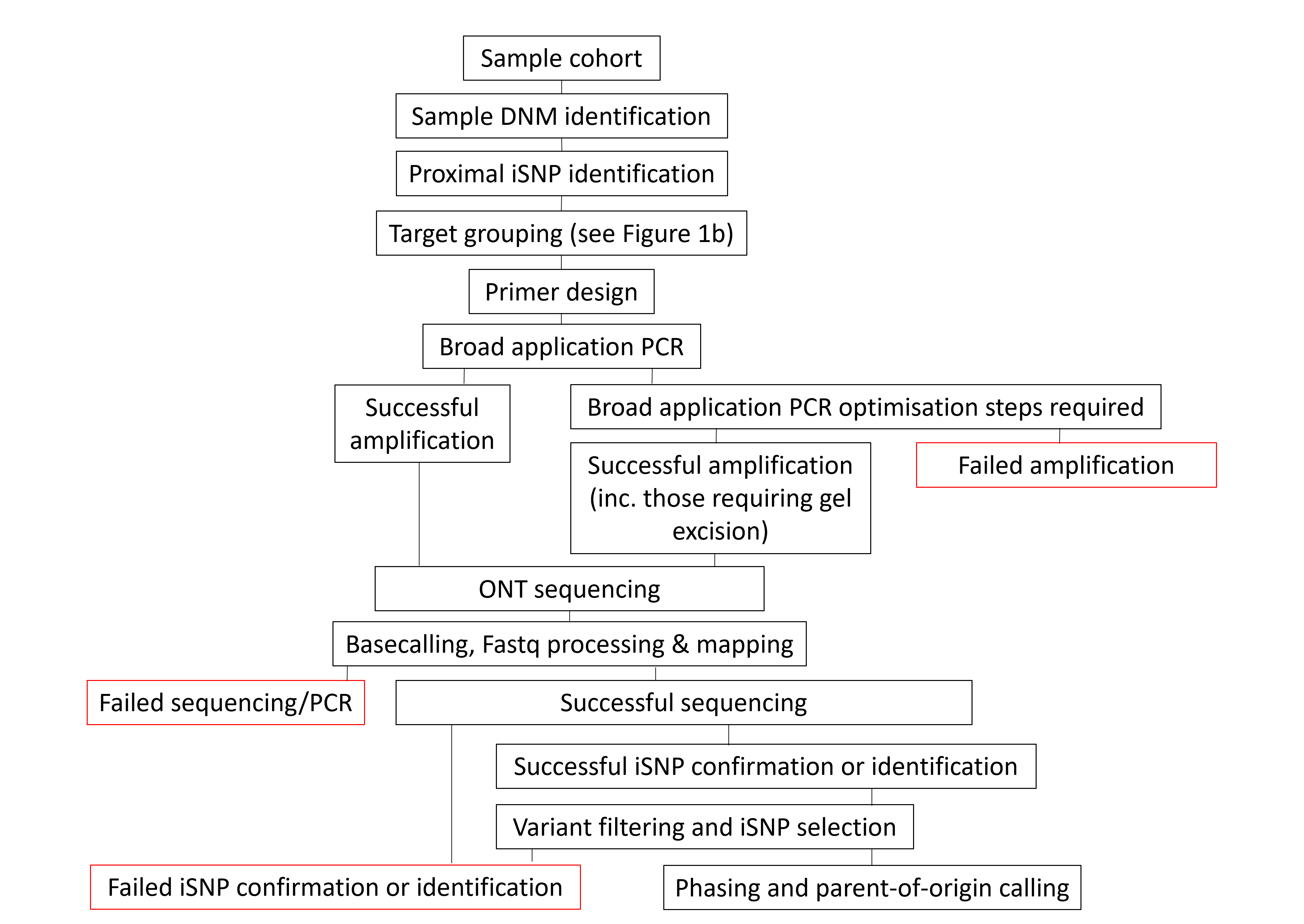

### Supplementary Figure 2

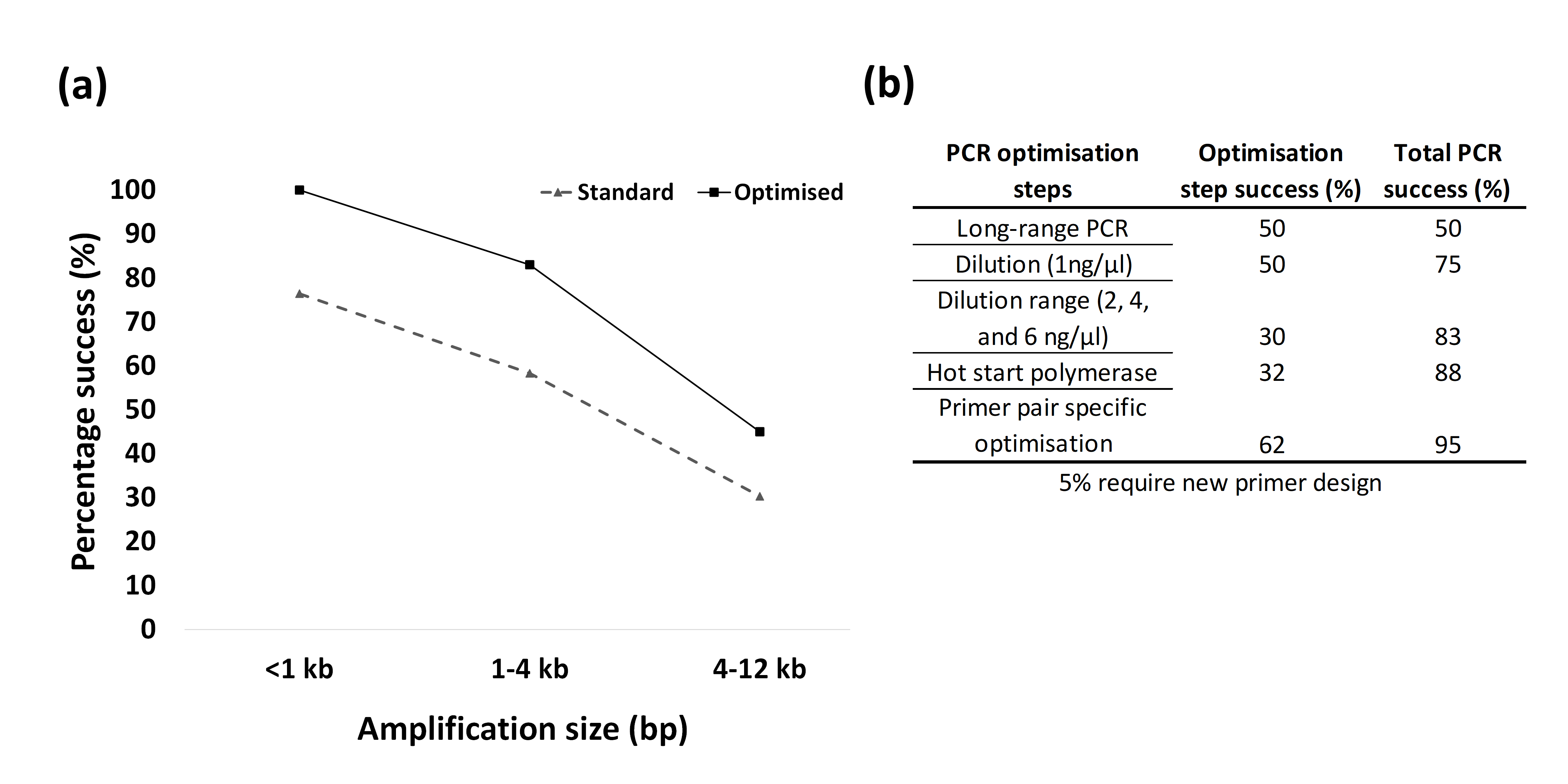

### Supplementary Figure 3

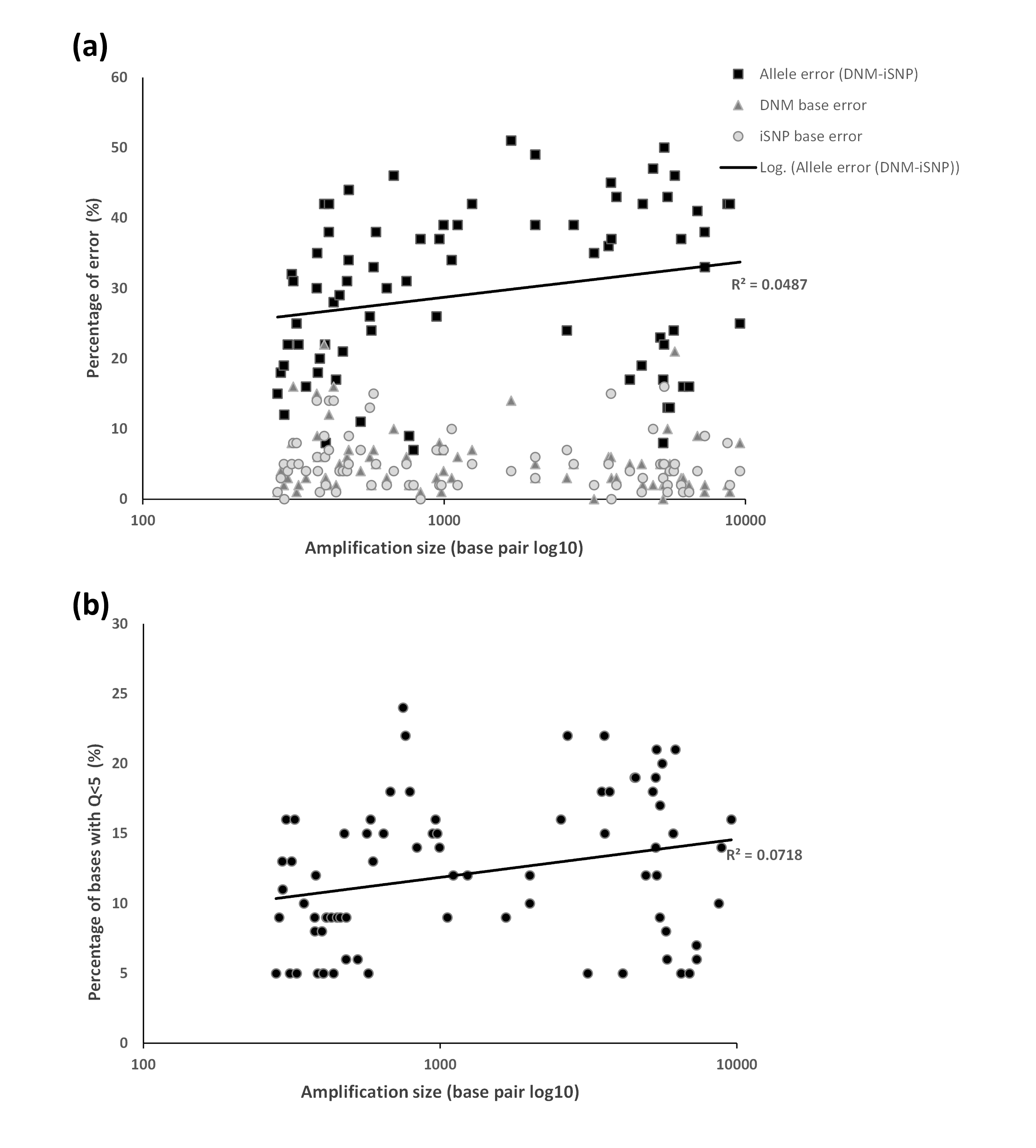

### Supplementary Figure 4

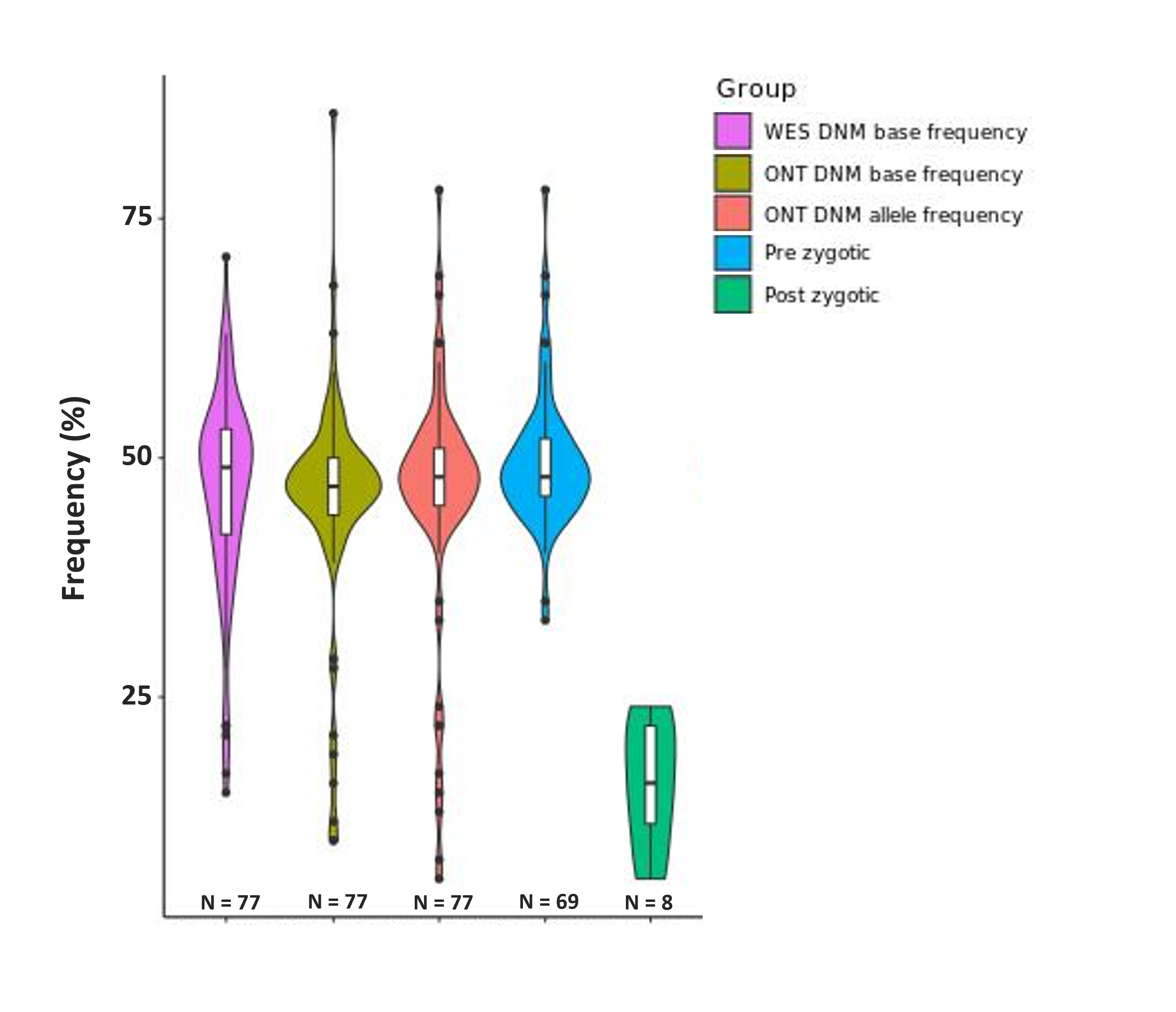

### Supplementary Figure 5

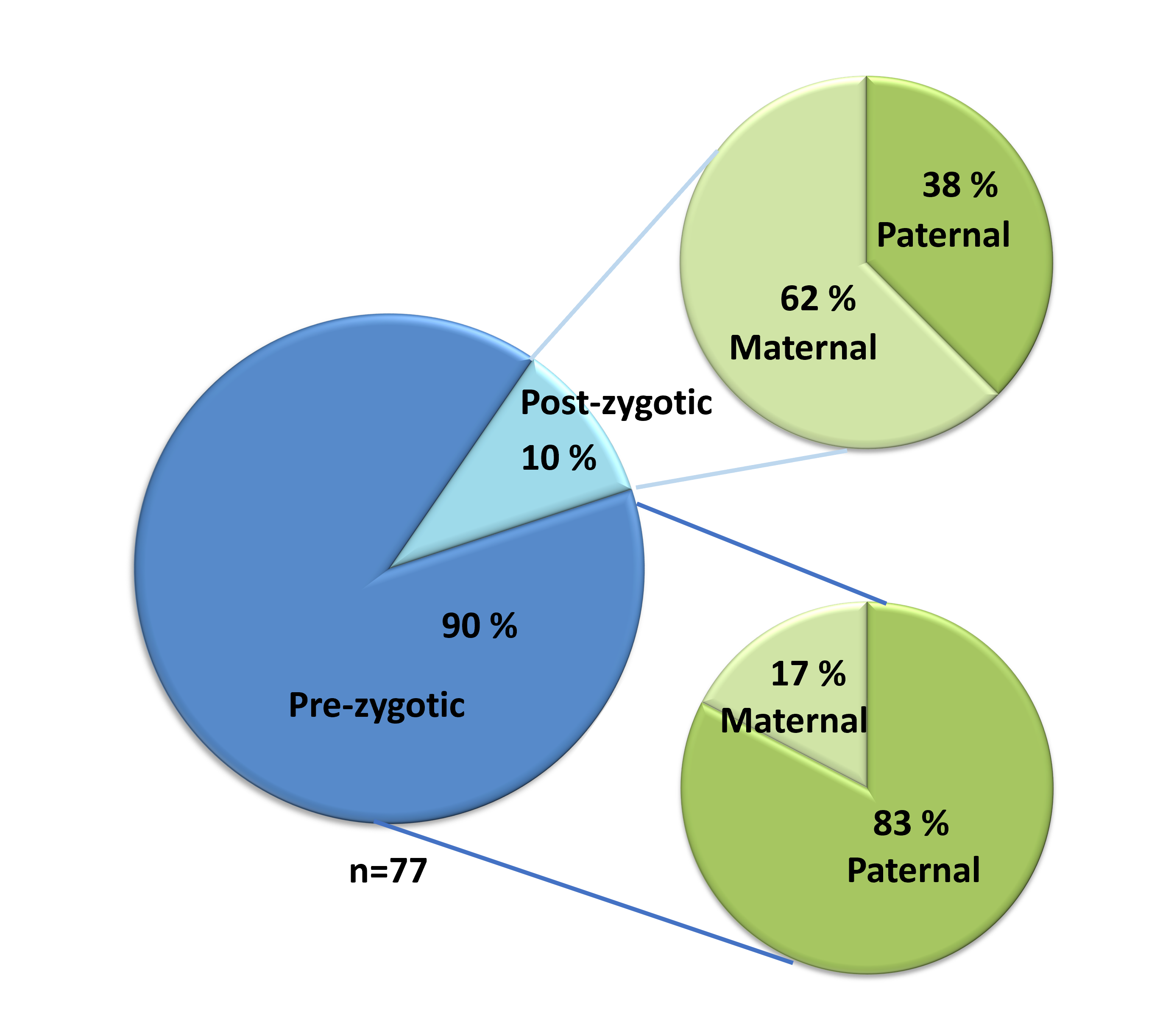

### Supplementary Figure 6

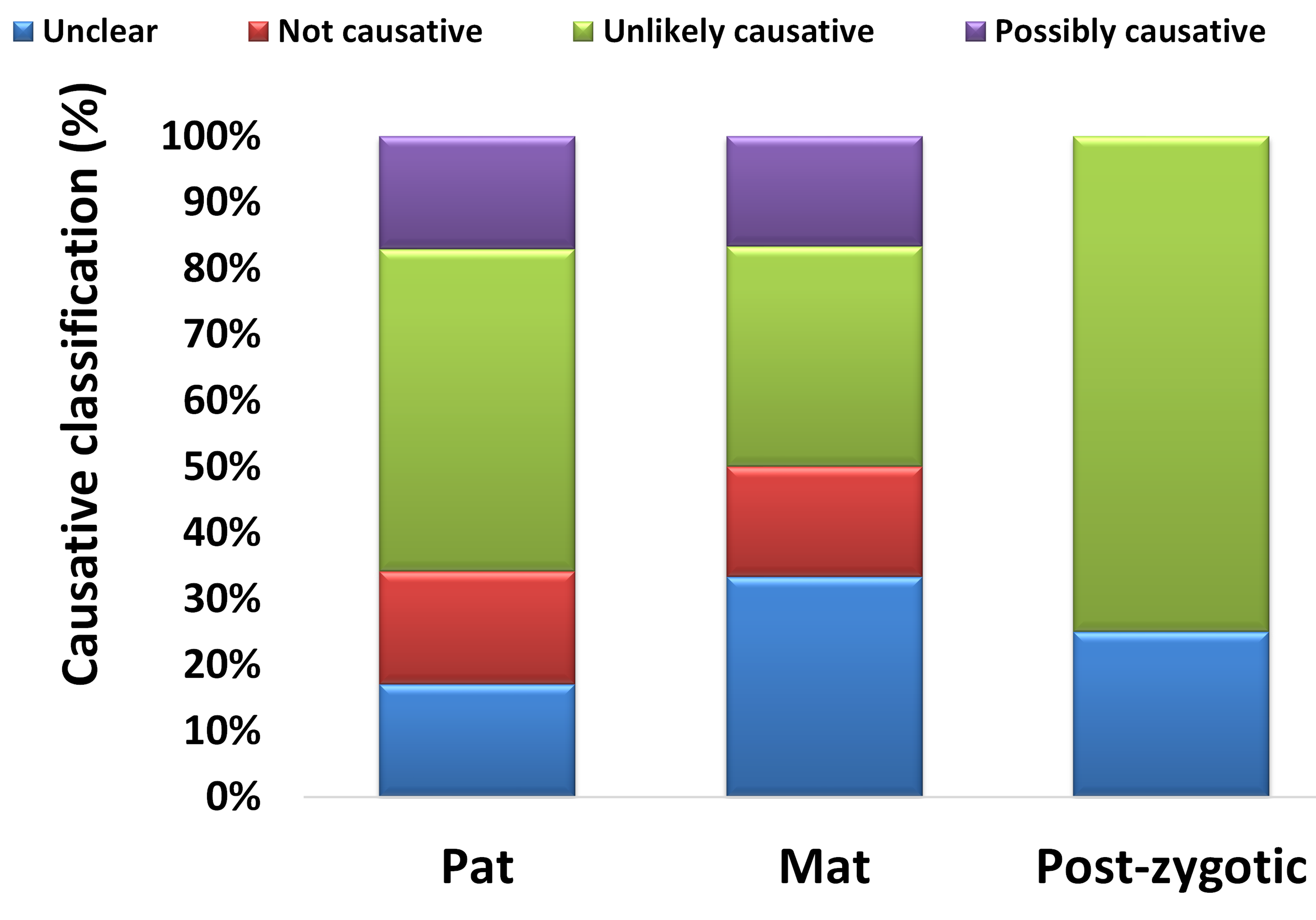

### Supplementary Figure 7

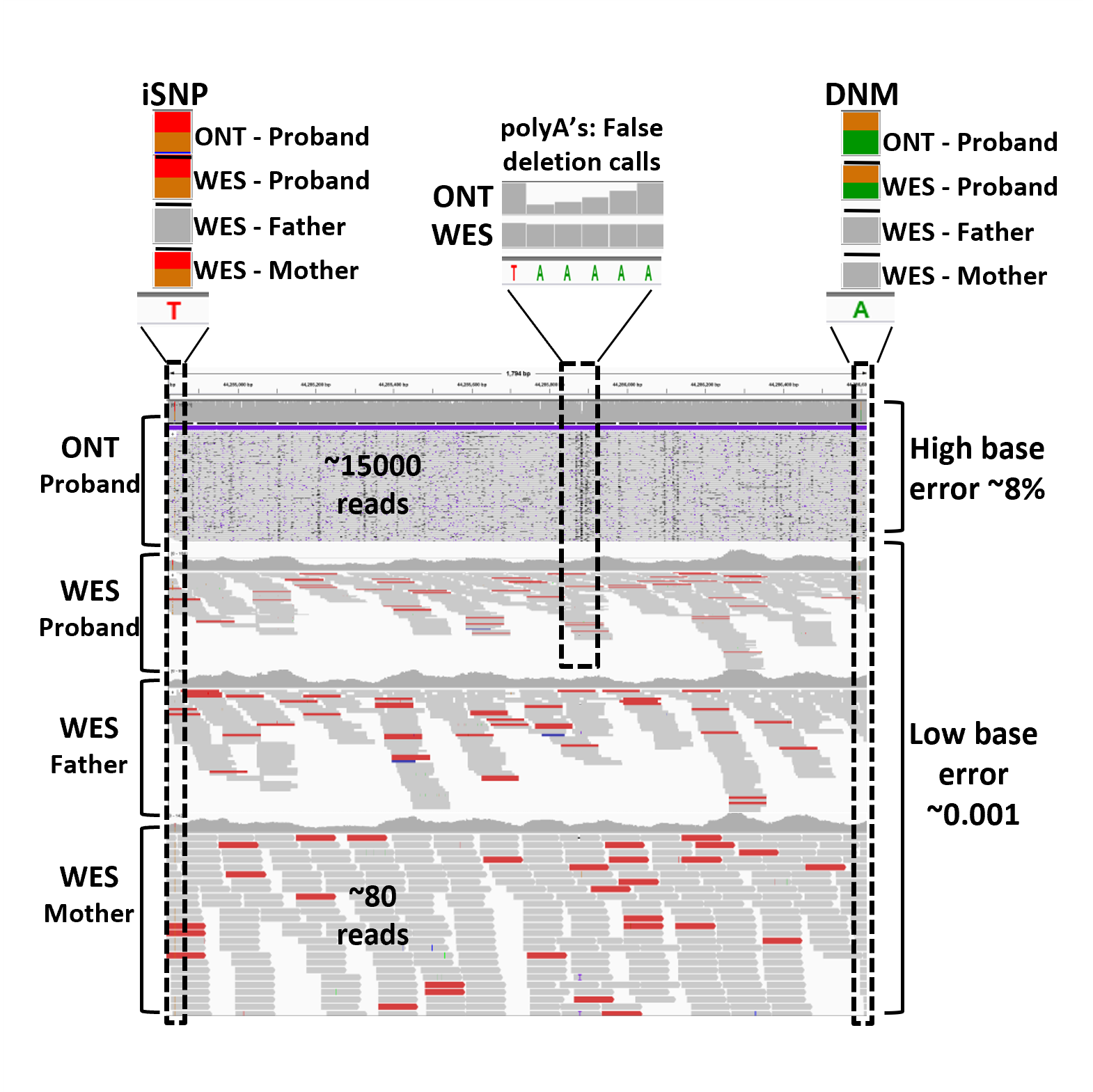
